# Super-multiplexed fluorescence microscopy via photostability contrast

**DOI:** 10.1101/258889

**Authors:** Antony Orth, Richik N. Ghosh, Emma Wilson, Timothy Doughney, Hannah Brown, Philipp Reineck, Jeremy Thompson, Brant C. Gibson

## Abstract

Many areas of biological research rely heavily on fluorescence microscopy to observe and quantify the inner workings of the cell. Traditionally, multiple types of cellular structures or biomolecules are visualized simultaneously with spectrally distinct fluorescent labels. A high degree of multiplexing is desirable as it affords the experiment greater information content, speeding up research timelines. Multiplexing can be increased by imaging a larger number of spectral channels, however, the wide emission spectra of most fluorophores limits multiplexing to four or five labels in standard fluorescence microscopes. Further multiplexing requires another dimension of contrast. Here, we show that photostability differences can be used to distinguish between fluorescent labels. By combining photobleaching characteristics with a novel unmixing algorithm, we resolve up to three fluorescent labels in a single spectral channel and unmix fluorescent labels with nearly identical emission spectra. We apply our technique to organic dyes, autofluorescent biomolecules and fluorescent proteins, and show that the latter are particularly well suited to our method as their bleaching is often reversible. Our approach has the potential to triple the multiplexing capabilities of any digital widefield or confocal fluorescence microscope with no additional hardware, making it readily accessible to a wide range of researchers.

## Introduction

Fluorescence microscopy is widely used in biological research for high contrast imaging with unrivalled specificity^1^. In a typical assay, 2-4 cellular targets are each labeled by fluorescent species with distinct spectral emission^2^. Combinations of different excitation sources and emission filters are then used to provide spectral contrast between the fluorescent species. In practice, there is inevitable mixing between fluorescent channels because fluorophores are excited and emit over a finite bandwidth. Fluorescent emission from a given fluorescent species contributes to the signal in multiple emission channels. This spectral cross-talk can be significant when using more than 3 fluorescent probes, necessitating unmixing algorithms for successful separation^3,4^. Generally, spectral unmixing procedures work well, however, one cannot isolate more fluorescent probes than one has spectral channels. High-end filter-based fluorescence microscopes have at most five excitation/emission filter combinations and can therefore spectrally separate up to 5 fluorescent species. Alternatively, spectrometer-based imaging systems can in principle acquire dozens of spectral channels, at the added cost of a large filter set or spectral imaging module. Unfortunately, even with a large number of spectral channels it is extremely challenging to identify more than 4-5 fluorescent species in a sample, regardless of the spectral measurement scheme^5,6^. This is because of the wide spectral width of each fluorophore (~40-50 nm with broad tails), the need for excitation windows (~20 nm minimum per excitation source) and the finite spectral bandwidth of the visible spectrum (300 nm). However, in many areas of biology there is a growing need to separate more objects or structures or to perform several fluorescence based sensing experiments simultaneously to increase the information content derived from fluorescence microscopy assays.

Here we introduce a method for expanding the number of fluorescent species that can be simultaneously multiplexed in an image, without resorting to large filter sets or spectral detectors. We show that one may discriminate between fluorescent species by leveraging a photophysical process inherent in all organic fluorophores – photobleaching. Instead of using spectral signatures to distinguish fluorescent species, we use their photobleaching behaviour as an identifying property. This technique, which we will call *bleaching-assisted multichannel microscopy* (BAMM), can be applied either by itself or in conjunction with spectral filters and is even capable of discriminating between fluorophores with nearly identical emission spectra.

Fluorescence photobleaching refers to a decrease in emission intensity of a fluorescent sample over time under illumination. This decrease is the result of chemical reactions between optically excited fluorescent molecules and the surrounding medium^7^. Bleached fluorophores are irreversibly “turned off” and are no longer able to emit light. The emission intensity (*I*) of an ensemble of fluorophores decreases exponentially over time (*t*) according to *I*(*t*) = *I*(0)*e^−kt^*. The bleaching constant *k* depends on a myriad of experimental and environmental parameters in addition to the electronic structure of the fluorophore itself^1^. For example, excitation power, excitation wavelength, oxygen concentration, and a fluorophore’s energy level structure all affect its bleaching rate^7^. This means that spectrally identical fluorophores can have very different bleaching rates.

A variety of microscopy techniques rely on photobleaching. For example, photo-imprint microscopy can increase resolution beyond the diffraction limit in both in-plane and axial dimensions^8,9^. Another super-resolution method called bleaching/blinking assisted localization microscopy (BALM) uses discrete photobleaching events to localize single molecules^10^. The resulting images are similar to that created by stochastic optical reconstruction microscopy (STORM) or photoactivated localization microscopy (PALM)^11,12^, but sample preparation is greatly simplified because of the universal occurrence of fluorescence photobleaching. Yet another technique, fluorescence recovery after photobleaching (FRAP), has become a standard tool for investigating diffusion kinetics in living cells^13,14^.

In the mid-1990s, photostability was briefly investigated as a contrast mechanism for multi-probe fluorescence microscopy^15,16^, but suffered due to a lack of sophisticated unmixing algorithms and inadequate computing power. We take advantage of subsequent developments in non-negative unmixing approaches^17,18^, and introduce a specialized non-negative matrix factorization algorithm for photobleaching data. Moreover, we show that multiplexing can be further increased by combining spectral and photostability dimensions, while 3D BAMM imaging can be achieved by employing the photorecovery characteristic common to many fluorescent proteins. Our work establishes photostability as a *bona fide* optical dimension for extended multiplexing without specialized sample preparation or additional microscope hardware.

## Results

### BAMM with fluorescent beads

In this section, we demonstrate the working principle of BAMM using a model system consisting of five different types of fluorescent beads (Spherotech and Sigma Aldrich, see *Methods*). The beads have peak emission wavelengths ranging from 500 - 700 nm, and are imaged simultaneously in yellow (570 - 620 nm) and red (663 - 738 nm) spectral channels using a confocal microscope (Fig. 1a). Bead types I-IV are all > 2μm in diameter and are therefore easily resolvable by the confocal microscope. However, bead type V is only 80nm in diameter, and therefore appears as a dim amorphous red background since the beads themselves are spatially unresolvable.

**Figure 1.**
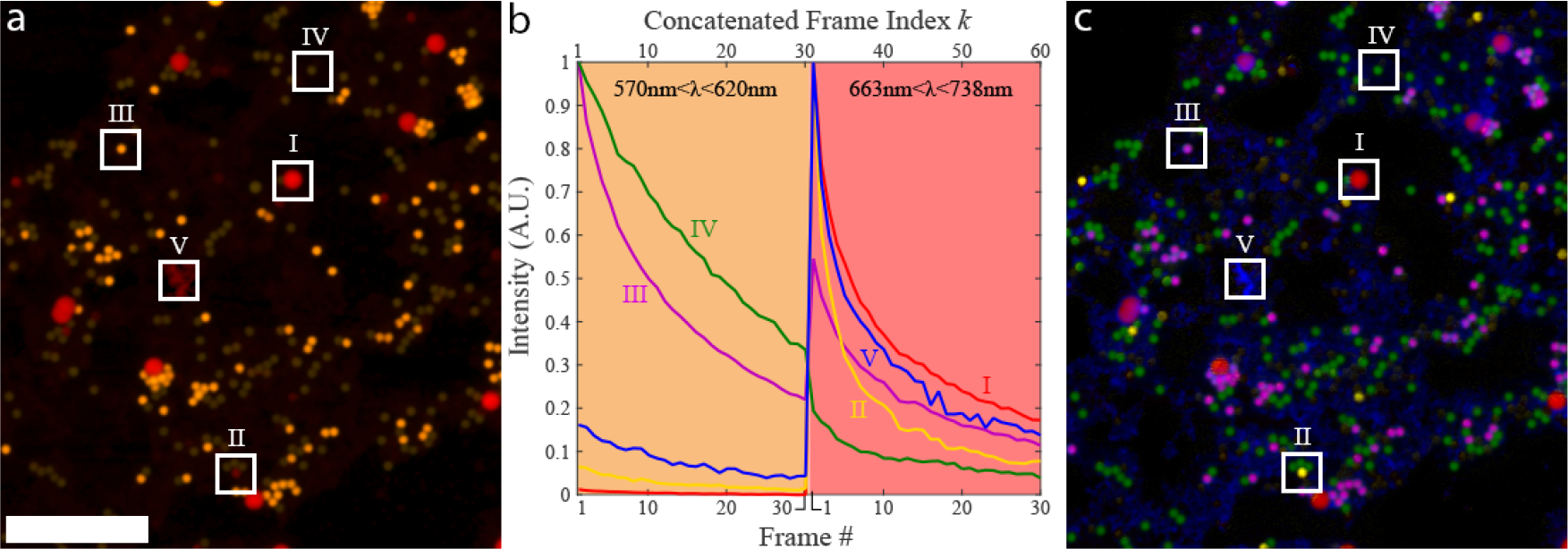
BAMM with beads, a) The first frame of a 30-frame photobleaching experiment, with each frame consisting of a yellow/red dual spectral channel image of a mixture of 5 different fluorescent beads. Examples of different bead types are boxed and labeled as I-V. The red channel is gamma corrected to enhance dim pixels (for display only). Boxes II, IV and V has increased brightness for visibility. Scale bar is 50 μm. b) Time traces of beads I-V in (a), during the photobleaching experiment. Concatenated frames # 1-30 correspond to frames 1-30 in the yellow channel (570-620 nm) of image (a). Concatenated frames #31-60 correspond to frames 1-30 from the red channel (663-738 nm) of image (a). Each bead type has a unique spectral-bleaching “fingerprint”, c) False-coloured unmixed image of all five bead types. Abundance maps for bead types I-V are coloured red, yellow, purple, green and blue, respectively. The unmixing fidelity of this process is shown in Fig. SI.

The first step in BAMM is to record a timelapse of sample bleaching. The sample is repeatedly imaged with one or multiple lasers simultaneously causing the sample to fade. We record a timelapse movie, followed by background subtraction and drift compensation pre-processing steps (see *Methods*)^19^. Assuming that several fluorescent species are present in a sample, they can all potentially contribute to the signal collected in a given pixel. In principle, determining the relative abundances of these fluorophore species via bleaching can be achieved by fitting the amplitudes of a multi-exponential decay at each pixel. This basic problem occurs in many arenas from magnetic resonance imaging^20^ to fluorescence lifetime imaging^21,22^ and nuclear physics^23^. There are a wide range of computational approaches to this challenge, such as maximum likelihood estimation^23^, the method of least-squares^24^, method of moments^25^ and the Gardner Transform^26^. However, our problem is more general as it includes both spectral and bleaching information, so we extract the spectral and photobleaching characteristics from the dataset itself. This self-calibrated approach avoids using physical models of photobleaching that may not be consistent with real world samples. We will refer to the spectral-bleaching characteristic of a fluorophore as its spectral-bleaching fingerprint. An analogous quantity has recently been used to unmix fluorescent probes based on their fluorescence lifetime and emission spectra^22^. In comparison, BAMM requires an order of magnitude less total integration time per pixel (~100μs vs. 1ms) in confocal microscopy, and can also be implemented in a widefield microscope, both requiring no additional hardware.

In order to obtain each species’ spectral-bleaching fingerprint, we manually identify 5 pixels in Fig. 1a, each of which contains signal from only one of each of the fluorophore types in the sample. At each pixel, we have an associated 30-frame bleaching curve in each of the two emission channels, for a total of 60 data points across spectral and temporal dimensions. The photobleaching traces for the two spectral channels are concatenated into the 60-frame spectral-bleaching fingerprint curves shown in Fig. 1b: concatenated frames 1-30 originate from the yellow spectral channel, concatenated frames 31-60 are from the red spectral channel. The abundance maps are subsequently obtained via standard non-negative least squares (NNLS) unmixing (See *Image Processing* – *Unmixing* in *Methods*)^17^. The result is a 5-channel image consisting of 5 independent fluorophore abundance maps (Fig. 1c), each of which indicates the relative concentration of given fluorescent species. Abundance maps for bead types I-V are colored red, yellow, purple, green and blue, respectively.

Cross-talk between abundance maps is shown in Supplementary Figure S1. Unmixing fidelity is generally high (<2.5% cross-talk) except for Type II beads (18.27% bleed through into Channel 1), due to the weak fluorescence of Type II beads. Though this sample contains mostly spatially non-overlapping objects, the mathematics employed here are fully capable of accounting for overlapping signals, as is shown in the following section. We note that using standard spectral unmixing techniques, it would only be possible to isolate *N* fluorescent species using *N* spectral channels. In Fig. 1, we improve on this limit by a factor of 2.5, unmixing 5 independent fluorophore abundance maps from two spectral channels. Spectral information, however, is not necessary for BAMM, which can separate spectrally identical fluorescent species provided their bleaching rate differs (Fig. S2).

### BAMM in cells

The unmixing methods used in BAMM are also capable of separating spatially overlapping objects often encountered in biological samples. To demonstrate this capability, we imaged muntjac skin fibroblasts labeled with Alexa Fluor 488 (actin) and Alexa Fluor 555 (mitochondria). A 473 nm laser excites both dyes and a single detector collects the dye emission over a spectral range that includes the peak emission wavelengths of both dyes (500-600 nm). The first frame of this monochromatic image series is shown in Fig. 2a. In general, overlapping structures preclude the possibility of finding pixels that contain each of the fluorophores in isolation. Thus, we cannot use the manual method of Fig. 1 for identifying the bleaching fingerprint of a fluorescent species. Instead, we use a non-negative matrix factorization (NMF) algorithm^18^ in MATLAB for simultaneous estimation of the bleaching behaviour of both dyes along with their spatial distribution (See *Image Processing* – *Unmixing* in *Methods)*. The non-negativity constraint helps to restrict possible solutions for the bleaching traces and the abundance maps to those with only positive values. This guarantees that the result is consistent with the fact that the intensity is a non-negative quantity.

**Figure 2.**
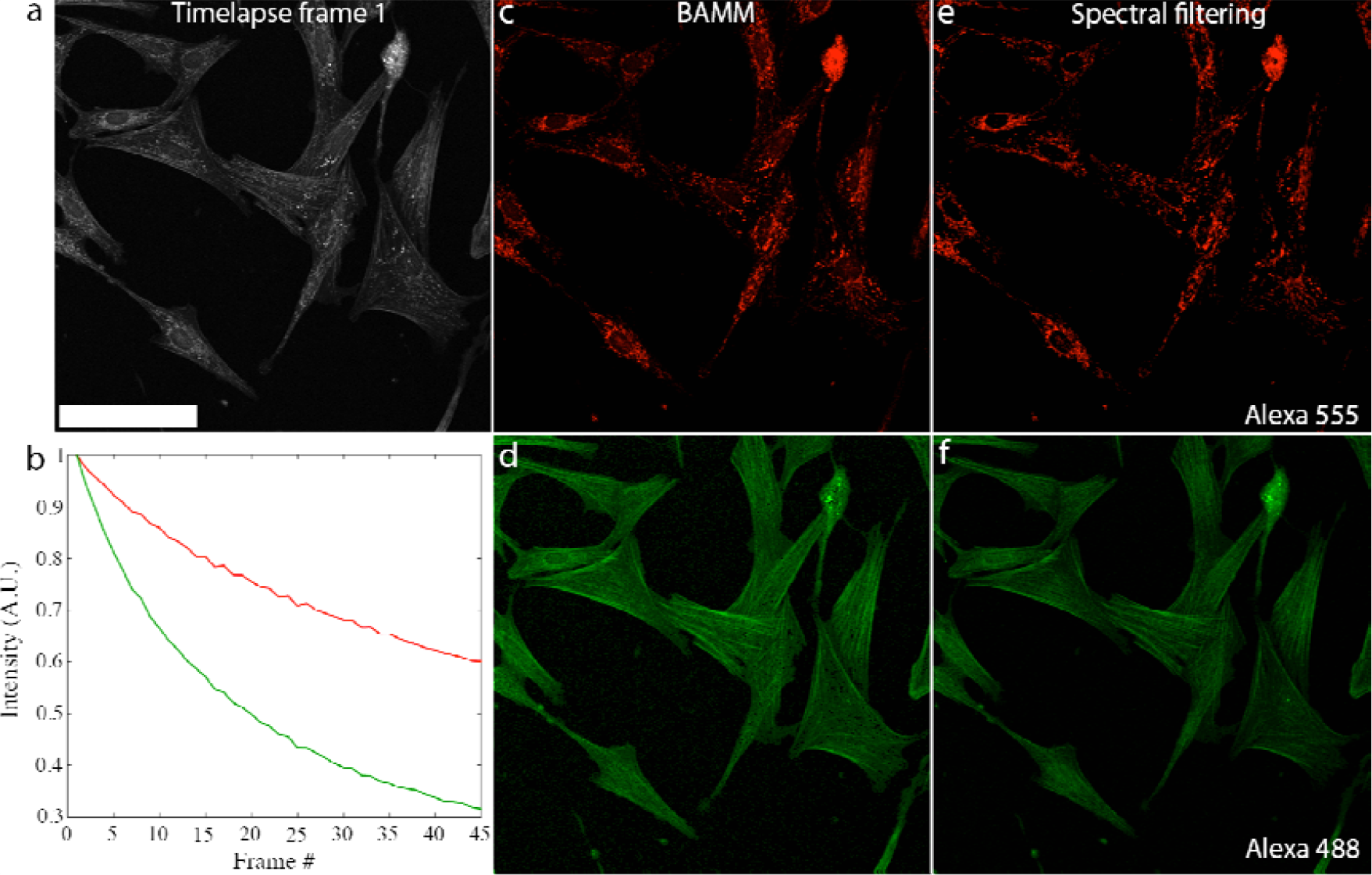
BAMM produces the correct abundance map with overlapping structures. a) First frame of a 45-frame bleaching timelapse of muntjac skin fibroblast cells under 473 nm illumination. A wide emission window is used (500-600 nm) to capture emission from both Alexa Fluor 488 and Alexa Fluor 555 dyes. Scale bar is 100 (μm. b) Bleaching traces of Alexa Fluor 555 (red) and Alexa Fluor 488 (green) as estimated by NMF. c) and d) Alexa Fluor 555 and Alexa Fluor 488 abundance maps extracted from the single-channel timelapse (a) using the estimated bleaching curves in (b) along with the entire 45-frame bleaching timelapse. Even though structures in both channels overlap spatially, they are unmixed successfully. e) Image of Alexa Fluor 555 distribution acquired using spectral filtering (570-670 emission window). f) Image of Alexa Fluor 488 distribution acquired using conventional spectral filtering (480-542 nm emission window). Both (e) and (f) were recorded before bleaching.

The NMF-estimated bleaching behaviours are shown in Fig. 2b: Alexa Fluor 555 (red) bleaches more slowly than Alexa Fluor 488 (green). The abundance maps of both dyes generated by NMF are shown in Figs. 2c and 2d (and in Supplementary Fig. S3c). Crucially, features are not duplicated between channels as occurs with incomplete unmixing. The spatial overlap between both fluorophore species is maintained. Reference images, acquired using spectral emission filters prior to bleaching (Figs. 2e and 2f) are nearly identical to those obtained using BAMM, demonstrating BAMM’s ability to produce high-quality multiplexed images of biological samples. The number of frames used dictates the quality of the unmixed images, with longer bleaching timelapse sequences with more frames and more bleaching yielding less noisy results at the expense of integration time (Supplementary Fig. S4).

Single-spectral-channel dual-labeling BAMM experiments can also be performed with fluorophores that are traditionally not separable using spectral filters. For BAMM, one particularly attractive application is the separation of spectrally overlapping traditional organic dyes from fluorescent proteins. These two types of emitters have very different photostabilities, and are thus readily unmixed using BAMM using either reference curves with NNLS estimation or NMF. We have used BAMM to separate Alexa Fluor 514 (tubulin) from GFP (mitochondria) in cells processed for Expansion Microscopy^27,28^ (Fig. 3a-c2, using reference curves) and Alexa Fluor 555 (Ki67) from RFP (Golgi) (Fig. 3d & e, via NMF) in HeLa cells. These samples were imaged with a widefield fluorescence microscope (Thermo Fisher Scientific CellInsight^TM^ CX7 HCA Platform), demonstrating that BAMM is compatible with both confocal and widefield microscopes as well as a super resolution technique in Expansion Microscopy. Using the latter we achieve an effective (pre-expansion) resolution of 163 nm using a 0.7 NA objective, as measured by the width of an isolated microtubule in the dual channel BAMM image. This resolution is slightly lower than the raw image data before BAMM processing (137 nm, see Supplementary Fig. S6), likely due to imperfect dedrifting correction.

**Figure 3.**
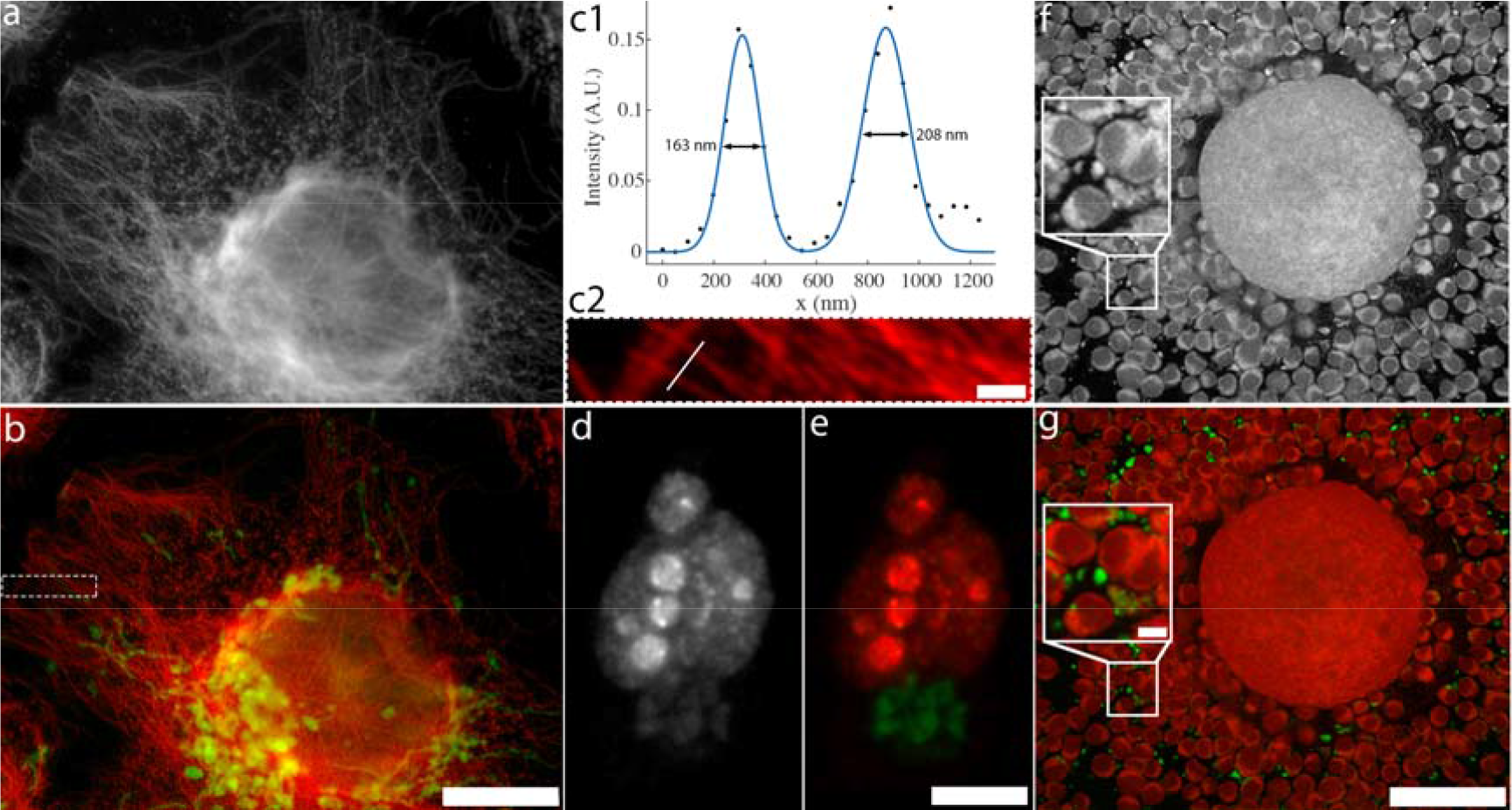
Dual-channel BAMM in cells. a) First frame of a single spectral channel bleaching timelapse of a U2OS cell processed for Expansion Microscopy, labeled with Alexa Fluor 514 (tubulin) and GFP (mitochondria). b) Dual-channel BAMM image of (a) using NNLS with reference bleaching curves (40 frames) for GFP and Alexa Fluor 514. Alexa Fluor 514 is colored red and GFP is colored green. Color histograms are modified for visibility of dim features. Dotted white border indicates region shown in (c2). Scale bar indicates 60 μm after expansion (13 μm before expansion). c1) Intensity trace along the white line in (c2), intersecting two isolated microtubules. The black dots indicate data points and the blue curve is a Gaussian fit to the intensity profile. The full width at half maximum of each peak is indicated on the plot. c2) Magnified version of the region within the dotted region in (b). The white line indicates the path of the intensity trace in (c1). GFP channel not shown for clarity. Scale bar is 5 μm after expansion (1.09 μm before expansion). d) First frame of a single spectral channel bleaching timelapse of a HeLa cell labeled with Alexa Fluor 555 (Ki67) and RFP (Golgi). e) Dual-channel BAMM image of (c) using NMF (45 frames). Alexa Fluor 555 is coloured red and RFP is colored green. Scale bar is 10 μm. f) First frame of a bleaching timelapse mouse cumulus-oocyte-complex autofluorescence. The boxed region is enlarged 2.5x in the inset. g) False-coloured BAMM image using NMF (50 frames). Scale bar is 50 μm, inset scale bar is 5 μm.

In addition to exogenous dyes and genetically engineered fluorescent proteins, many biomolecules are naturally autofluorescent. These molecules are not inherently designed for photostability or multichannel imaging, and therefore have highly overlapping spectral profiles and display a wide range of bleaching behaviour. The former makes them particularly hard to unmix spectrally and the latter is appealing for BAMM. Figure 3f is an autofluorescence image of a mouse cumulus-oocyte-complex (COC) within a wide yellow-to-red emission window (575-675 nm). Analysis with NMF reveals two distinct fluorescent populations with different photostability. In the unmixed image (Fig. 3g), photostable foci (green) can be seen amongst the less photostable autofluorescence of the oocyte and cumulus cells (red). This result suggests that photostability may be a valuable, yet overlooked form of fluorophore identification, particularly in spectrally crowded samples.

Unmixing three types of fluorophores via bleaching requires *a priori* information beyond non-negativity in order to reduce the solution space. To this end, we modify MATLAB’s alternating least squares NMF algorithm to force the solution to the bleaching curves to be monotonically non-increasing (see *Image Processing - Non-increasing NMF* in *Methods*). This guarantees that the NMF solution is physically consistent with the knowledge that fluorescent intensity must decrease over time. We call this approach the non-increasing NMF algorithm (NI-NMF). This added restriction of monotonically decreasing basis functions (the bleaching trace estimates) is crucial for 3-component unmixing using BAMM. Figure 4a shows a HeLa cell with 3 fluorescent emitters in a single spectral channel, while the 3-pseudo-color NI-NMF BAMM image is shown in Fig. 4b. This unmixing result represents a 3-fold increase over the traditional multiplexing limit with only one spectral channel. The associated estimated bleaching traces for this dataset are shown in Fig. 4c, for both NMF and NI-NMF unmixing. The NI-NMF algorithm correctly returns bleaching traces that only decrease over time whereas the unmodified NMF algorithm estimates bleaching behavior that increases over certain time intervals, leading to incomplete and physically inconsistent unmixing results.

**Figure 4.**
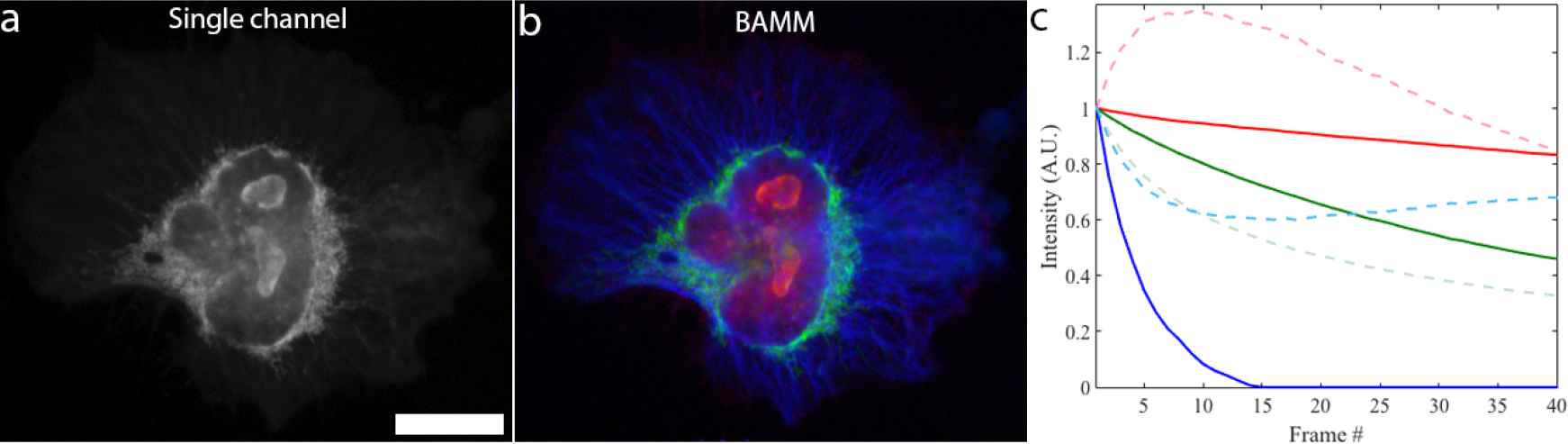
BAMM unmixing of 3 fluorescent labels from a single spectral channel. a) The first frame (of 40) of the BAMM bleaching timelapse, using 485nm LED excitation. This LED excites Alexa Fluor 555 (Ki67), GFP (mitochondria) and Alexa Fluor 430 (microtubules). Scale bar is 20 μm. b) False-color BAMM image obtained via NI-NMF, showing the fluorescent emitter distributions. Alexa Fluor 555 in red, GFP in green and Alexa Fluor 430 in blue. c) Bleaching traces for each fluorophore type estimated by NI-NMF (solid) and NMF (dotted). Solid curve colors match the color scheme in (b).

### Reverse BAMM

Fluorescent proteins are well known to undergo reversible photobleaching and photoswitching^29,30^. This is an inherent property of the fluorophore and is not to be confused with diffusion recovery processes used in techniques such as FRAP^13^. Reversible photobleaching presents an additional means of photochemical contrast: fluorophores with different photobleaching recovery behaviours can be unmixed using BAMM. Previous work demonstrated that fluorescent proteins can be distinguished based on photoswitching properties alone, provided that the photoswitching properties of each fluorescent protein in the system is complementary^31^. We show that this approach can be modified for cases where one of the fluorescent species displays only irreversible bleaching – eliminating the requirement for specialized combinations of fluorescent proteins with opposite photoswitching behaviours.

As a simple example of this concept we demonstrate unmixing of GFP from Alexa Fluor 488 in fixed CaCO2 cells by sequentially bleaching with 473nm excitation and then inducing photobleaching recovery with a 405nm beam. This procedure modulates the intensity of GFP, while Alexa Fluor 488 undergoes slight monotonic bleaching (Fig. 5a-d). Because of this large contrast, unmixed images can be obtain either from pairs of bleached/recovered images (frames 10-11, Fig. 5e) or from the bleaching portion of the intensity trace (frames 11-20, Fig. 5f). Here, the reversible photobleaching properties of GFP significantly reduces the number frames needed for unmixing. This enables 3D BAMM imaging (Supplementary Movie 1) and may enable BAMM imaging of live cells in future studies.

**Figure 5.**
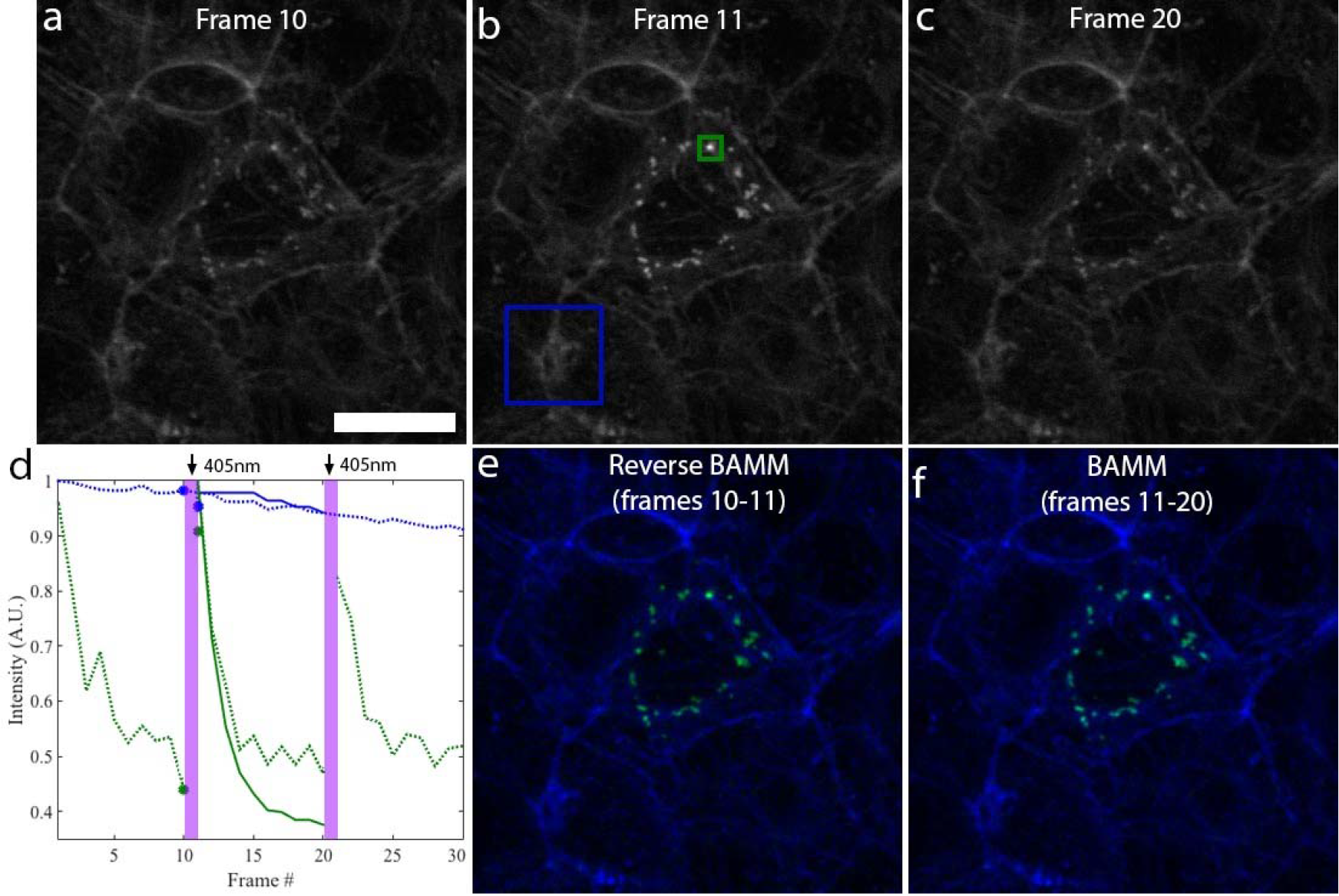
Unmixing with reversible photobleaching. a)–c) The 10^th^, 11^th^ and 20^th^ frame of a reversible bleaching timelapse of GFP (Golgi) expressing CaC02 cells, stained with Alexa Fluor 488 (actin). The sample is irradiated with 405nm light between the 10^th^ and 11^th^ frame, inducing photorecovery of GFP. The sample is then imaged with 473nm light from frames 11-20, causing a decrease in signal over the entire sample. Scale bar is 25 μm. d) The intensity trace of the central pixel in the green square in (b) is shown in the dotted green curve; the mean value of all pixels within the blue square is shown in the dotted blue curve. Both curves are normalized so that their maximum value = 1. Estimated (NI-NMF) bleaching traces between frames 11-20 are shown in the solid blue and green curves for Alexa Fluor 488 and GFP, respectively. Blue and green points denote the estimated (NMF) photorecovery intensity points for Alexa Fluor 488 and GFP, respectively. Vertical purple bars indicate 405nm laser irradiation for photorecovery, e) Unmixed image of Alexa Fluor 488 (blue) and GFP (green) using NMF on frames 10 and 11. f) Unmixed image of Alexa Fluor 488 and GFP using NI-NMF on frames 11-20.

## Discussion

We have shown that BAMM is compatible with both laser scanning confocal and widefield microscopes. Widefield illumination has the advantage that all fluorophores in a weakly absorbing 3D sample receive the same peak excitation power. In a confocal microscope this is not the case – the excitation intensity is highest in the middle of the focal spot. Fluorophores above and below the focal plane accumulate the same total dose during image acquisition, but experience a smaller peak excitation power. If nonlinear photobleaching effects are large, this can cause a height dependent photobleaching response. However, this effect is mitigated by the confocal aperture, which blocks most out–of-plane emission from reaching the detector. We did not observe any depth dependent bleaching rates, provided we used a pinhole size smaller than or equal to the Airy disc.

An even illumination profile is critical for BAMM because the bleaching rate is a function of excitation intensity. Two species at different locations in the field-of-view can experience the same photobleaching rate if they are subject to different excitation intensities. If the illumination profile is known *a priori*, this effect could be mitigated by adjusting the temporal binning at each pixel to compensate for the spatially dependent excitation dose. A more straightforward approach is to simply crop the field of view so that the illumination profile is uniform within a smaller region.

To investigate whether the intrinsic bleaching rate of fluorophores varies throughout fixed cell samples, we fit a single exponential model at each pixel of single spectral channel images of muntjac skin fibroblasts (Supplementary Fig. S5). We observe no significant systematic spatial variation in bleaching rates of either Alexa Fluor 488 or Alexa Fluor 555 within cells; the bleach rate variation is largely within the 95% confidence bounds of the fitting parameter estimates. This is perhaps not surprising since fixed cell samples are subject to chemical treatment to prevent degradation and to allow fluorophores to pass through certain cellular structures. The chemical environment, which affects the bleaching rate, is therefore likely quite homogeneous, as opposed to the live cell case. Furthermore, we note that concentration can, in principle, affect photostability via dye-dye interactions^7^, however, this only occurs when fluorophores are so concentrated that they can directly interact with one another. In this regime, the linearity between fluorophore concentration and emission is undermined via dye self-quenching^1^, making regular fluorescence quantitation impossible. This concentration regime is avoided by necessity for any quantitative experiment. As such, the bleaching rate is likely to remain independent of concentration for any experiment where fluorescence intensity is linear with concentration, which is the case for nearly all properly conducted experiments.

We have shown that BAMM is compatible with Expansion Microscopy^27,28^, a relatively new route towards super resolution microscopy. We expect that BAMM will also be suitable for use with structured illumination, so long as accurate de-drifting correction is applied. Compatibility with fundamentally nonlinear techniques such as two photon^32^ and stimulated emission depletion is less clear as they are more susceptible to nonlinear bleaching^32^. Single molecule localization microscopy (SMLM) techniques such as stochastic optical reconstruction microscopy^11^ are fundamentally different because they identify single molecules. On this scale, bleaching is a discrete process. Instead of calculating abundances at each pixel, an SMLM approach to BAMM might instead classify a fluorophore based on how many photons it emitted before bleaching (or the photobleaching recovery rate for fluorescent proteins). This could not only simplify multi-label SMLM but also allow more efficient photon collection and lower system cost by reducing the number of filters and light sources needed.

In summary, we have introduced a novel microscopy method, BAMM, which uses photobleaching as a contrast mechanism. Though photobleaching is usually thought of as being detrimental to fluorescence imaging, here we harness this ubiquitous effect to multiplex up to three fluorescent labels into a single spectral channel. BAMM reports on a fundamentally different property of fluorophores (their photostability) than standard spectral filter based fluorescence microscopy. As such, BAMM can be used to distinguish multiple fluorescent species given only a single emission channel, and separate nearly spectrally identical fluorophores. Reverse BAMM takes advantage of reversible photobleaching, which provides yet another means of photochemical contrast. We suspect that monitoring the rate of photobleaching recovery in addition to regular irreversible photobleaching may yet provide a means to even further multiplexing, with potential applicability to live cell imaging. Our work complements other recent reports of fluorescence super-multiplexing methods where fluorescent labels are distinguished based on properties beyond standard excitation and emission spectra, such as fluorescence lifetime^22^ and stimulated Raman scattering^33^. Unlike these techniques, however, BAMM does not require any additional hardware and can be performed on a standard digital fluorescence microscope – widefield or confocal.

## Methods

### Materials

#### Beads

Bead types I-IV were sourced from Spherotech. Type I: Spherotech “Sky Blue”; type II: Spherotech “Blue”; type III: “Purple”; type IV: “Yellow”. Bead type V: Chromeon 642 from Sigma Aldrich.

#### Muntjac skin fibroblasts

This sample (Figs. 2 and S3-S5) is a commercially available microscope slide: Thermo Fisher Scientific FluoCells prepared slide #6 (F36925). The sample is labeled with Alexa Fluor 488 Phalloidin (F-actin), Alexa Fluor 555 (mitochondria) and TO-PRO-3 (nucleus). We only use Alexa Fluor 488 and Alexa Fluor 555 due to restrictions on the maximum bandwidth of the emission filter on the Olympus FV1200 microscope.

### Preparation of adherent cell samples

#### HeLa cell culture

A culture of human cervical adenocarcinoma cells HeLa, CCL-2™ (ATCC, Manassa, VA, USA) was maintained using the ATCC standard culture method for this cell line. Briefly, the cells were incubated at 37°C with 5% CO_2_ in Eagle’s Minimum Essential Eagle Medium (MEM, Gibco™, USA) containing 1% penicillin-streptomycin, 1% sodium pyruvate, 1% non-essential amino acids, and 10% fetal bovine serum (FBS, Thermo Fisher Scientific, USA). Subculture was performed when cells were 80% confluent by using 0.25% Trypsin (Thermo Fisher Scientific, USA) for 5-10 minutes at 37°C. For specific fluorescent labelling low passage number HeLa cells were seeded into the wells of sterile 96-well plates with cover glass bottoms (Nunc™, Thermo Fisher Scientific, catalog # 164588) at a density of 5 x 10^3^ cells/well, incubated overnight, fluorescently labeled and then imaged.

#### U20S cell culture

A culture of human osteosarcoma cells U2OS, HTB-96™ (ATCC, Manassa, VA, USA) was maintained using the ATCC standard culture method for this cell line. Briefly, the cells were incubated at 37°C with 5% CO2 in McCoy’s 5A Medium (Gibco™, USA) containing 1% penicillin-streptomycin, and 10% fetal bovine serum (FBS, Thermo Fisher Scientific, USA). Subculture was performed when cells were 80% confluent by using 0.25% Trypsin (Thermo Fisher Scientific, USA) for 5-10 minutes at 37°C. For labeling and processing the cells for Expansion Microscopy, low passage number U2OS cells were seeded into the individual wells of a sterile 8-well CultureWell™ multiwell chambered coverslip (Thermo Fisher Scientific, catalog # C24779) at a density of ~ 10^4^ cells/well, incubated overnight, fluorescently labeled and then imaged.

#### Fluorescent labeling of HeLa and U2OS cells

Labeling of the Golgi apparatus, mitochondria, or nuclei were done using the commercially available transfection reagents CellLight® Golgi-RFP BacMam 2.0, CellLight® Mitochondria-GFP BacMam 2.0, or CellLight® Nucleus-CFP BacMam 2.0 respectively (catalog #s C10593, C10600, and C10616, Thermo Fisher Scientific, USA). Briefly, 20, 30, or 50 viral particles/cell of the respective CellLight reagents were added to each well according to the manufacturer’s directions at the same time as cell plating, followed by 24-48 hours incubation. Upon successful expression of the fluorescent proteins, the cells were fixed by incubation at room temperature for 15 minutes with 10% formalin neutral buffered solution (Sigma-Aldrich, USA), then washed three times with phosphate buffered saline pH7 (PBS, Thermo Fisher Scientific, USA), and then permeabilized with 0.1% TritonX-100 in PBS (Thermo Fisher Scientific, USA). The Ki67 and microtubules were labeled by indirect immunofluorescence as follows: Cells were incubated at room temperature for 1 hour with either a rabbit monoclonal antibody against Ki67 (clone D2H10, catalog # 9027, Cell Signaling Technology, USA), a mouse monoclonal antibody against alpha-tubulin (clone DM1A, catalog # MS-581-P1, Thermo Fisher Scientific, USA), or both antibodies together, diluted at 1:500 in PBS. The cells were then washed three times with PBS containing 1% bovine serum albumin (BSA, catalog # 37525, Thermo Fisher Scientific, USA) before incubation at room temperature for 1 hour with the appropriate fluorescent secondary antibody diluted 1:200 in PBS (all secondary antibodies were from Thermo Fisher Scientific, USA). The secondary antibody used to identify the Ki67 was Alexa Fluor® 555 goat anti-rabbit IgG (catalog # A21429); the secondary antibodies used to identify the alpha-tubulin were either Alexa Fluor® 430 goat anti-mouse IgG (catalog # A11063), Alexa Fluor® 514 goat anti-mouse IgG (catalog # A31555), or Alexa Fluor® 555 goat anti-mouse IgG (catalog # A21424). The cells were then washed three times with PBS before imaging.

#### Caco-2 cells

A culture of human colon colorectal cells Caco-2, HTB-37™ (ATCC, Manassa, VA, USA) was maintained using the ATCC standard culture method for this cell line. Briefly, the cells were incubated at 37°C with 5% CO_2_ in Dulbecco’s Modified Eagle Media (DMEM, Gibco™, USA) containing 100 U/mL penicillin-streptomycin and 10% fetal bovine serum (FBS, Sigma-Aldrich, USA). Subculture was performed when cells were 80% confluent by using 0.25% Trypsin (Sigma-Aldrich, USA) for 5-10 minutes at 37°C and centrifugation at 200 × g at 4°C for 5 minutes.

For specific fluorescent labelling low passage number Caco-2 cells were seeded onto a sterile cover glass at a density of 2 × 10^5^ cells and incubated overnight (19 hours) as above. Labeling of the Golgi apparatus was done using the commercially available transfection reagent CellLight® Golgi-GFP, BacMam 2.0 (Life Technologies, USA). Briefly, 2 µL per 50,000 cells was incubated with the above cells for 48 hours. Upon successful expression of the GFP, the cells were fixed by incubation at room temperature (19°C±1) for 10 minutes with 10% formalin neutral buffered solution (Sigma-Aldrich, USA). The fixed cells were washed three times with phosphate buffered saline pH7 (PBS, Gibco™, USA). The cells were permeabilized using 0.2% TritonX-100 in PBS (Sigma-Aldrich, USA). The actin filaments (Factin) were stained using 110 nM AlexaFluor488–Phalloidin in 200 µL of PBS for 30 minutes at room temperature with gentle mixing. The cells were then washed three times with PBS before mounting in PBS solution containing 10% Glycerol (Sigma-Aldrich, USA).

#### Expansion Microscopy

U2OS cells were fluorescently labeled as described above, and then processed for Expansion Microscopy as previously described^28^. Briefly, PBS was removed from the fluorescently labeled cells in 8-well CultureWell™ multiwell chambered coverslips, and 35 µL of 0.1 mg/mL Acryloyl X (catalog # A20770, Thermo Fisher Scientific) was added to each well. After incubating the cells in Acryloyl X for 2 hours at room temperature, the cells were washed three times with PBS, and then followed by the gelation, digestion, and expansion steps. The monomer gelation solution was prepared on ice; to make 9.4 mL monomer solution, 1 mL of 10x PBS and 0.9 mL water were combined with the appropriate volumes of the following components stock concentrations to have a final concentration of: 86 mg/mL sodium acrylate, 25 mg/mL acrylamide, 1.5 mg/mL N,N⍰-Methylenebisacrylamide, 110.7 mg/mL sodium chloride (Sigma-Aldrich, USA). The accelerator (tetramethylethylenediamine; TEMED) and initiator (ammonium persulfate; APS) were from 10% stocks (Sigma-Aldrich, USA) and were added to the monomer solution to final concentration of 0.2%, and then the combined gelation mixture was rapidly added to wells of the chambered coverslip and placed in a humidified 37°C incubator for 1 hour. For digestion, the gels were then fully immersed in Proteinase K (8 units/mL, Thermo Fisher Scientific) in 50 mM Tris (pH 8), 1 mM EDTA, 0.5% NaCl, 0.5% Triton X-100 (Sigma-Aldrich, USA) and incubated at room temperature overnight. The individual digested gels were then transferred to the wells of a coverglass-bottomed 6-well plate (catalog # NC0452316, Fisher Scientific). Excess volume of doubly deionized water was added to each well for 15-60 min, and this step was repeated 3-4 times until the size of the expanding gel plateaued. The samples were imaged using a 20× 0.7 numerical aperture air objective.

To determine the amount of expansion, images of the CellLight CFP labeled nuclei were imaged before and after the cells had been processed for Expansion Microscopy, and their areas were compared. The square root of the ratio of the post-expansion nuclear area to the pre-expansion nuclear area gives the amount of linear expansion, which in this case was found to be ~4.6, consistent with previous reports^27,28^.

To determine the effective resolution of the expanded samples, we measured the intensity profiles across 10 individual fluorescently labeled microtubule fibers (Fig. S6). We then applied a .Gaussian fit to each intensity profile, which yielded a mean FWHM of 137 nm, with standard deviation of 14 nm. This figure is below the optical diffraction limit with visible light and is consistent with previous Expansion Microscopy results^27,28^.

### COC sample preparation

#### Animals

All experiments were approved by the University of Adelaide Animal Ethics Committee and were conducted in accordance with the Australian Code of Practice for the Care and Use of Animals for Scientific Purposes. All mice were maintained in 14L:10D photoperiod conditions and given water and rodent chow *ad libitum*.

#### Media

Unless otherwise specified, all reagents and antibodies were purchased from Sigma-Aldrich (St. Louis, MO). All media used were as previously described^34^. Briefly, COCs were liberated into αMEM supplemented with bovine serum albumin (BSA; ICPbio, Glenfield, New Zealand). Recombinant human follicle-stimulating hormone (50 mIU/ml; Organon, Oss, The Netherlands) was subsequently used for IVM. Maturation media was pre-equilibrated for at least 4 h prior to use at 37°C in a humidified 6% CO_2_ atmosphere.

#### COC collection and IVM

Female mice were administered (i.p.) 5 IU equine chorionic gonadotropin (eCG; Folligon, Intervet, Boxmeer, The Netherlands). 46 h post-eCG injection, ovaries were collected and COCs liberated from antral follicles. COCs were then placed in maturation medium and matured for 17 h in a volume of 50 μl medium/COC at 37°C under paraffin oil, in humidified air comprised of 20%0_2_,6% CO_2_ and N_2_balance. Following maturation, COCs were fixed in 4% paraformaldehyde in phosphate buffered saline (PBS) and mounted on glass slides using DAKO Fluorescence Mounting Medium (Dako, NSW, Australia).

### Data acquisition

#### Confocal microscopy

Data from Fig. 1 and Supplementary Fig. S2 were acquired using a Nikon AR1 confocal microscope, equipped with continuous wave (CW) excitation lasers at λ=405, 488, 561 and 640 nm. Images were acquired using a 40x, 0.95 NA microscope objective, with a pixel dwell time of 1 µs and 2x line averaging. The data in Fig. 1a are colour coded with yellow and red channels corresponding to emission windows 570-620 nm and 663-738 nm, respectively. All emission channels are recorded simultaneously, and all 4 lasers excite the sample simultaneously with laser powers of (0.20, 1.10, 1.20, 0.09) mW for (405, 488, 561, 640) nm lasers, respectively. In Supplementary Figure S2a, the data is from the 663-738 nm channel. The sample was excited with the 640 nm laser at a power of 0.4 mW. For all data acquired with the Nikon AR1 confocal microscope, the confocal pinhole was set to 80% of the diffraction limit at 640 nm.

Data from Figs. 2 and 3f was acquired using an Olympus FV1200 spectral confocal microscope equipped with a 20×0.75NA air (Fig. 2), and 60x 1.3 NA silicone immersion (Fig. 3f) objective lenses.

For Fig. 2, the sample was excited with a 473nm CW laser with a dwell time of 8 μs, with a power of 0.27 mW. The emission was collected over a single emission window stretching from 500-600 nm. The confocal aperture was set to the diffraction limit for 473 nm.

For Fig. 3f the sample was excited with 473 and 559 nm CW lasers simultaneously (power 0.14 mW for each laser) with a pixel dwell time of 2 μs. The emission was collected within a range of 575-675 nm, using a variable bandpass filter built into the microscope. The confocal aperture was set to 60% of the diffraction limit at 559 nm.

Data for Fig. 5 was also acquired using a FV1200 confocal microscope, with 473 nm excitation 0.032 mW, using a 20x 0.75NA air objective. Dwell time was 2 µs, emission window from 485-545nm, confocal aperture set to the diffraction limit at 473nm. Photorecovery was induced by rastering a 0.070 mW 405nm focal spot twice over the field of view with 4 µs dwell time per pixel.

For Supplementary Movie 1, the sample volume is first imaged with a 473nm laser beam on an Olympus FV1200 confocal microscope. Subsequently, a second volume is imaged with a 405nm activation scan before imaging with 473nm for each slice. The activation scan reverses GFP photobleaching at each z-height, creating contrast between the first (no photobleaching reversal) and second (photobleaching reversal) volumes. Imaging parameters are the same as for Fig. 5, except a 1.3 NA 60x silicone immersion objective was used. The spacing between confocal planes was 310 nm.

#### Widefield Microscopy

Data from Figs. 3a-d and 4 were obtained from a Thermo Fisher Scientific CellInsight^™^ CX7 High Content Analysis (HCA) platform. Imaging parameters were as follows.

Figs. 3a,b & S6: 20×/0.7 NA objective, with 227 nm pixel size in image space. Excitation provided by a 485nm LED excitation with a 20nm FWHM bandpass filter. Fluorescence emission was detected through a 27nm FWHM bandpass filter centered at 542nm. Integration time 6.0s/frame.

Fig. 3d&e: 40×/0.75NA objective, with 2×2 binning for 227nm pixel size in image space. Sample was sequentially illuminated and imaged with 386nm, 485, 549 and 560nm LEDs for exposure times of 3.0s, 3.0s, 1.0s and 1.0s, respectively. Software autofocus was performed before each iteration of the LED illumination sequence with 386nm excitation. Bleaching timelapse processed for BAMM used only information from 549nm excitation images, which were collected through a 5-band bandpass filter (transmission bands of the 5-band emission filter are (center wavelength/FWHM bandpass): 438/47 nm, 521/22 nm, 604/30 nm, 704/54 nm, and 810/85 nm).

Fig. 4: 40×/0.75NA objective, with 2×2 binning for 227nm pixel size in image space. Sample was sequentially illuminated and imaged with 386nm, 485, 549 and 560nm LEDs for exposure times of 1.5s, 2.0s, 0.75s and 0.75s, respectively. Software autofocus was performed before each iteration of the LED illumination sequence with 386nm excitation. Bleaching timelapse processed for BAMM used only information from 485nm excitation images, which were collected through the 5-b and bandpass filter as above.

#### Fluorescence spectra

Spectra in Supplementary Fig. S2c were obtained by drying each type of bead onto a separate slide, exciting at 520 nm and recording the emission while scanning with a homebuilt confocal microscope equipped with a Pixis 100 spectrometer (Princeton Instruments).

### Image processing

#### Timelapse pre-processing

Before unmixing, raw data are background-subtracted by manually selecting a blank area of the image, and subtracting the mean pixel value over this region from the image at each time point. This eliminates signal arising from autofluorescence bleaching from the substrate and negates detector background noise drift throughout the experiment. The data are then corrected for sample drift. The spatial offset vector of each frame in the dataset is calculated via cross correlation with the first frame. Each frame is offset back an amount equal to the negative offset vector.

#### Unmixing

For Figs. 1, 3b and Supplementary Fig. S2 we use the MATLAB NNLS function *lsqnonneg* to solve the following system of *T* equations and *N* unknowns^17^

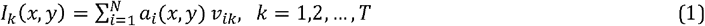

Where *I_k_* (*x*, *y*) is the (measured) intensity at pixel (*x*, *y*) for concatenated frame index *k, a_i_(x*, *y*) ≥ 0 is the relative scalar abundance of fluorophore species *i* at pixel *(x*, y) and *v_ik_* is the *k*^th^ entry of the *T*-element (spectral-)bleaching fingerprint for fluorophore type *i*. Note that the system is overdetermined when *T > N*. The extra information in this overdetermined system is crucial for noise suppression (see Supplementary Figure S3). The non-negativity prior for *a_i_* prevents the unmixed abundances from reaching nonphysical negative values and improves unmixing fidelity, though its implementation requires an iterative approach. The *i*^th^ image - the abundance map of fluorophore type *I -* is given by *a_i_*(*x, y*) and ideally contains only signal from the *i*^th^ fluorescent species.

For Fig. 3b, reference bleaching curves obtained with singly-labeled samples under the same imaging conditions are used.

For Figs. 2, 3e and 5, the bleaching fingerprint of each fluorophore is estimated by the MATLAB non-negative matrix factorization function *nnmf*. This algorithm attempts to find non-negative bleaching traces and fluorophore abundances that solve the mixing problem of Eq. 1 in the least squared sense, using the alternating least squares (ALS) procedure^18^. We supply the *nnmf* function with the principal components of photobleaching curves as the initial solution estimate (see Fig. S3a). For added robustness, we take the optimal solution out of three replicates – the first seeded with principal components as initial solutions and the following two with random initial guesses. Typical results required no more than 25 iterations.

For Supplemental Movie 1, unmixing is achieved by running the NMF procedure on a slice that contains both GFP and Alexa Fluor 488 (slice 9/14), yielding estimates of the magnitude of bleaching of Alexa Fluor 488 (bleaching recovery for GFP) from volume 1 to 2. These estimates are then used to unmix the remainder of the slices in the volume.

#### Non-increasing NMF

For Fig. 4, we implement a modification to the standard ALS NMF procedure, which we call non-increasing NMF (NI-NMF). At each iteration, if the value of a bleaching trace estimate B(t) at time t=t_n+ 1_ exceeds the value of the bleaching trace estimate at time t_n_, then we set B(t_n+1_)=B(t_n_). This is modification is performed for each time point in succession so that the value of the bleaching trace is monotonically decreasing. We run NI-NMF for 25 replicates, with the first one being seeded with the principal components, and select the result with the lowest mean-squared error. For the NMF-estimated bleaching traces in Fig. 4c, we run the identical procedure, but without the non-increasing constraint. For all NI-NMF and NMF procedures it was found to be beneficial to exclude dim pixels to reduce the influence of noise and decrease computation time. We typically exclude all pixels with less brightness less than 1-10% of the brightest pixel in the dataset. Once the estimate for the bleaching curves is found, the least squares solution for the abundances at every pixel in the image is found by using the timelapse data and the estimated bleaching curves together with the MATLAB backslash operator (a QR solver), and then setting negative abundances to zero. This is equivalent to including all pixels on the final iteration of the ALS (NI-) NMF algorithm.

## Data Availability

The datasets generated during and/or analysed during the current study are available from the corresponding author on reasonable request.

## Acknowledgements

This work was supported by the ARC Centre of Excellence for Nanoscale BioPhotonics (CE140100003). B. C. G. acknowledges the support of an ARC Future Fellowship (FT110100225). We would like to thank the staff at the RMIT MicroNano Research Facility for access to microscopy equipment. We would also like to thank Calum Drummond for the use of the Olympus confocal microscope used in this work.

## Author Contributions

A.O. conceived the experiments. A.O, B.G. and R.N.G. designed the experiments. H.B., J.T., R.N.G and E.W. developed biological samples. A.O., R.N.G. and T.D. performed the experiments and analyzed data. All authors contributed to writing the paper.

## Competing Financial Interests

The authors declare no competing financial interests.

